# The non-dominant AAA+ ring in the ClpAP protease functions as an anti-stalling motor to accelerate protein unfolding and translocation

**DOI:** 10.1101/585554

**Authors:** Hema Chandra Kotamarthi, Robert. T. Sauer, Tania. A. Baker

## Abstract

ATP-powered unfoldases containing D1 and D2 AAA+ rings play important roles in protein homeostasis, but uncertainty about the function of each ring remains. Here we use single-molecule optical-tweezers to assay mechanical unfolding and translocation by a variant of the ClpAP protease containing an ATPase-inactive D1 ring. This variant displays substantial mechanical defects both in unfolding and translocation of protein substrates. Notably, when D1 is hydrolytically inactive, ClpAP often stalls for times as long as minutes, and the substrate can “back-slip” through the enzyme when ATP concentrations are low. The inactive D1 variant also has substantially more difficulty traveling in the N-to-C direction on a polypeptide track than moving C-to-N. These results indicate that D1 normally functions as an auxiliary/regulatory motor to promote uninterrupted enzyme advancement that is fueled largely by the D2 ring.

## Introduction

AAA+ enzymes (<u>A</u>TPases <u>a</u>ssociated with various cellular <u>a</u>ctivities) perform mechanical processes in cells by operating as tiny motors that convert the chemical energy of ATP hydrolysis into work that typically involves macromolecular remodeling^1, 2^. In all domains of life, a subfamily of AAA+ enzymes promotes protein homeostasis by degrading damaged or misfolded proteins^3, 4^. AAA+ proteases contain a barrel-shaped peptidase and a hexameric AAA+ unfoldase with an axial pore that can bind specific peptide tags (degrons) in target proteins. Following binding, cycles of ATP hydrolysis in the AAA+ hexamer power substrate unfolding and subsequent translocation into the peptidase chamber for degradation^3^. For example, the *Escherichia coli* ClpAP protease, which degrades ssrA-tagged proteins and other substrates, consists of the AAA+ ClpA hexamer and the ClpP peptidase^5–7^.

ClpA hexamers contain 12 active sites for ATP hydrolysis, six in each of two distinct AAA+ rings; a similar double-ring architecture is present in the hexamers of related enzymes (ClpB, Hsp104, Cdc48/p97, and NSF) that function without proteolytic partners in protein remodeling. ClpA in the absence of ClpP also has remodeling activity^8^. Why ClpA and other double-ring enzymes need two rings is unclear. Some mutagenesis experiments suggest that one ring performs most of the ATP hydrolysis and functions as the dominant enzyme motor. It has been proposed that the second ring assists in the nucleotide-stabilized assembly of the hexamer^9–11^, but single-ring AAA+ enzymes also assemble into stable hexamers, and there would appear to be many simpler ways to stabilize the double-ring enzymes.

The top ClpA AAA+ ring, where substrates first enter the axial pore, is called D1, whereas the bottom ring, which is proximal to ClpP, is called D2 (Fig. 1a)^12, 13^. ATP hydrolysis in one ring or the other can be eliminated by mutating a either of the conserved glutamate in the D1 Walker-B motif (E286) or in the D2 Walker-B motif (E565). Studies of these enzyme variants show: (*i*) D2 appears to catalyze most ATP hydrolysis, although elimination of hydrolysis in D1 can reduce ATPase activity by as much as 2-fold in the presence of some protein substrates; (*ii*) inactivating D1 generally has modest effects on substrate degradation, whereas inactivating D2 impairs degradation severely; and (*iii*) ATP hydrolysis in both rings is important for degradation of substrates with very stable local structures^14^. Because the D2 ring appears to be most important for ATP-fueled degradation, we focused on determining how ATP hydrolysis in the less-understood D1 ring contributes to mechanical substrate unfolding and polypeptide translocation, critical steps in the overall degradation reaction.

**Figure 1:**
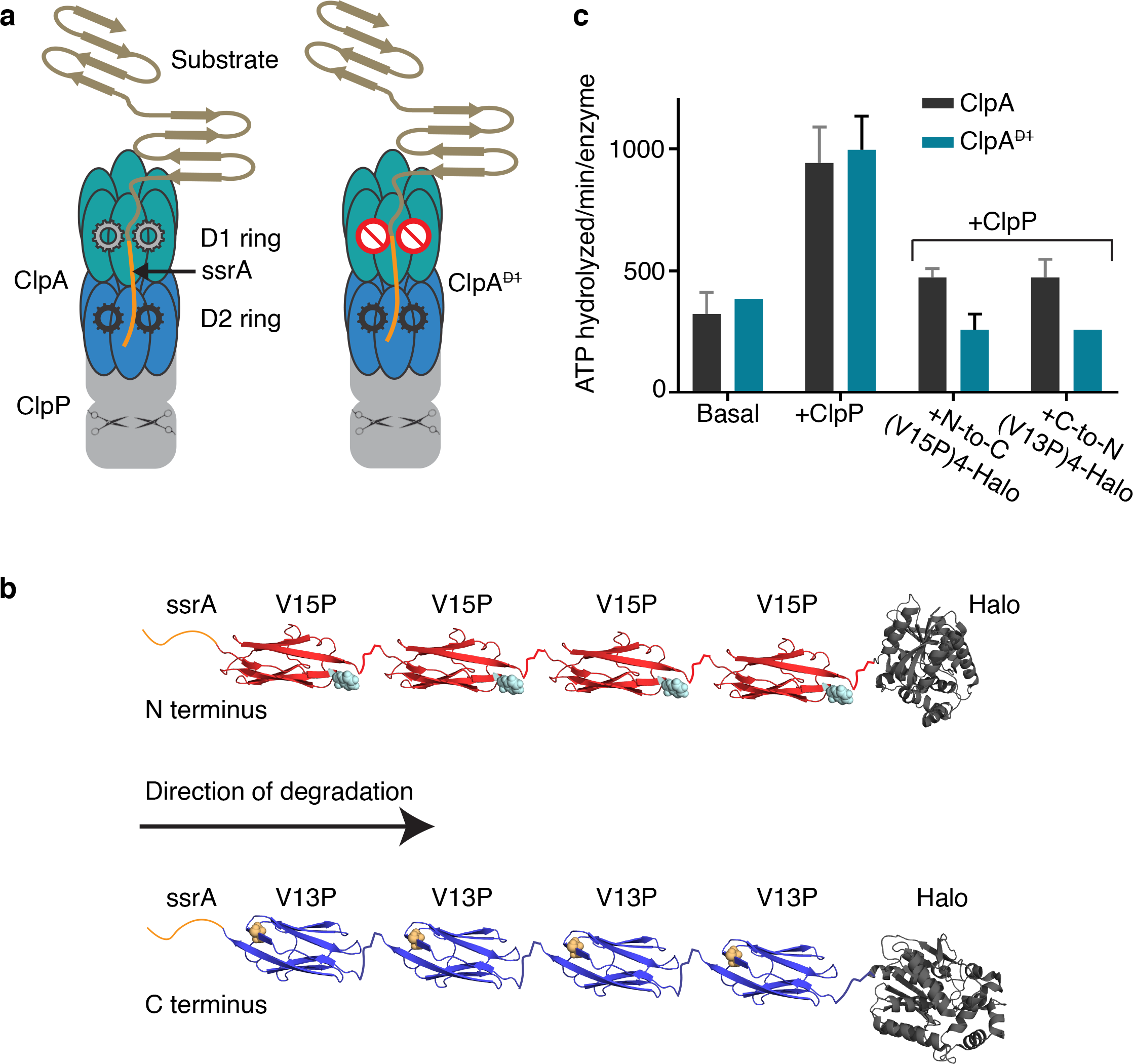
ClpAP protease, multidomain substrates, and ATP hydrolysis. (**a**) **Cartoon of ClpAP and ClpA^D1^P.** Each ClpA subunit contains two AAA+ ATPase modules that form the D1 (teal) and the D2 (blue) rings in the assembled hexamer. ClpP (gray) is a barrel-shaped serine peptidase; scissors schematically represent peptidase active sites. A model protein substrate engaged for unfolding in the ClpA pore is shown. ClpA^D1^ (right) contains a 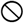 symbol in the D1 ring to indicate an ATP-hydrolysis defect. (**b) Schematic of ssrA-tagged multidomain substrates.** The ssrA tag (orange) is present at the N-terminus of ssrA-(V15P)_4_-Halo (red, top) or the C-terminus of Halo-(V13P)_4_-ssrA (blue). The sites of the V15P and V13P mutations in the titin I27 domain are represented as spheres (note the titin domains are in opposite orientations). The Halo domain, used for covalent linkage to DNA for optical trapping, is dark gray. (**c**) **Steady-state rates of ATP hydrolysis.** Rates for ClpA or ClpA^D1^ (0.1 μM hexamer) were measured in the absence of ClpP and substrate (basal), in the presence of ClpP (0.2 μM tetradecamer), or in presence of ClpP (0.2 μM tetradecamer) and substrate (5 μM). Values are averages ± 1 SD of three independent measurements.

Here, we use single-molecule optical-tweezer assays to investigate how ClpAP unfolding and translocation are affected when the D1 ring cannot hydrolyze ATP as a consequence of an E286Q mutation. We find that this D1-defective ClpAP (ClpA^D1^P) displays a marked increase in the frequency of pausing/stalling during translocation as well as substantial defects in unfolding activity. Hence, ATP hydrolysis in the D1 ring increases enzyme productivity by stimulating both of these key components of mechanical degradation. Surprisingly, the ClpA^D1^P defects were magnified when degradation proceeded in a N-to-C direction compared to C-to-N direction. Our results also support a role for D1 in maintaining substrate grip and preventing slipping.

## Results

### Single-molecule degradation by ClpA^D1^P

Here we used two multidomain substrates previously employed to analyze single-molecule degradation by wild-type ClpAP^15, 16^. Degradation was initiated either from the N-terminus or the C-terminus of these substrates by changing the position of the ssrA tag^15, 16^, which targets the protein to ClpA. For N-to-C degradation, the substrate contained an N-terminal ssrA tag, four human titin I27 domains with the mildly destabilizing V15P mutation, and a Halo domain (Fig. 1b, top). For C-to-N degradation, we initially tested a substrate with four V15P domains, Halo, and a C-terminal ssrA tag, but ClpA^D1^P degraded this substrate poorly. As a consequence, we used a substrate with four titin domains containing the more destabilizing V13P mutation, Halo, and an ssrA tag (Fig. 1b, bottom).

When protein substrate was absent, ClpA^D1^ hydrolyzed ATP at essentially the same rate as ClpA, and ClpP stimulated hydrolysis to similar extents for both enzymes (Fig. 1c). In the presence of either protein substrate, however, ClpA^D1^P had only 50-60% of the wild-type hydrolysis activity (Fig. 1c). Hence, when engaged with these substrates, the D1 ring of ClpA appears to contribute substantially, either directly or indirectly, to overall ATP-hydrolysis activity. Similar results were reported previously, although only certain substrates reduced ATPase activity for ClpA^D1^P in comparison with ClpAP^14^.

To probe the contribution of the D1 ring to unfolding and translocation by ClpA, we performed single-molecule optical-tweezer assays (Fig. 2a)^17, 18^. Biotinylated ClpA^D1^P was immobilized on a 1.25 μm streptavidin-coated polystyrene bead and the substrate Halo domain was attached, via a biotinylated DNA linker, to a 1.09 μm bead (Fig. 2a)^17, 19^. The beads were trapped by two laser beams and experiments were performed in a passive force-clamp mode at forces ranging from 8 to18 pN. After the two beads were tethered together via ClpA^D1^P on one bead binding to the substrate on the other, we monitored inter-bead distance as a function of time. Each sharp increase in distance reflects unfolding of a single domain, the following gradual decreases in distance are caused by translocation of the now unfolded polypeptide, and periods of no movement just prior to unfolding are the pre-unfolding dwell times. We also collected a handful of new traces of single-molecule degradation by ClpAP, but largely relied on our previously published analyses for comparisons^15, 16^.

**Figure 2:**
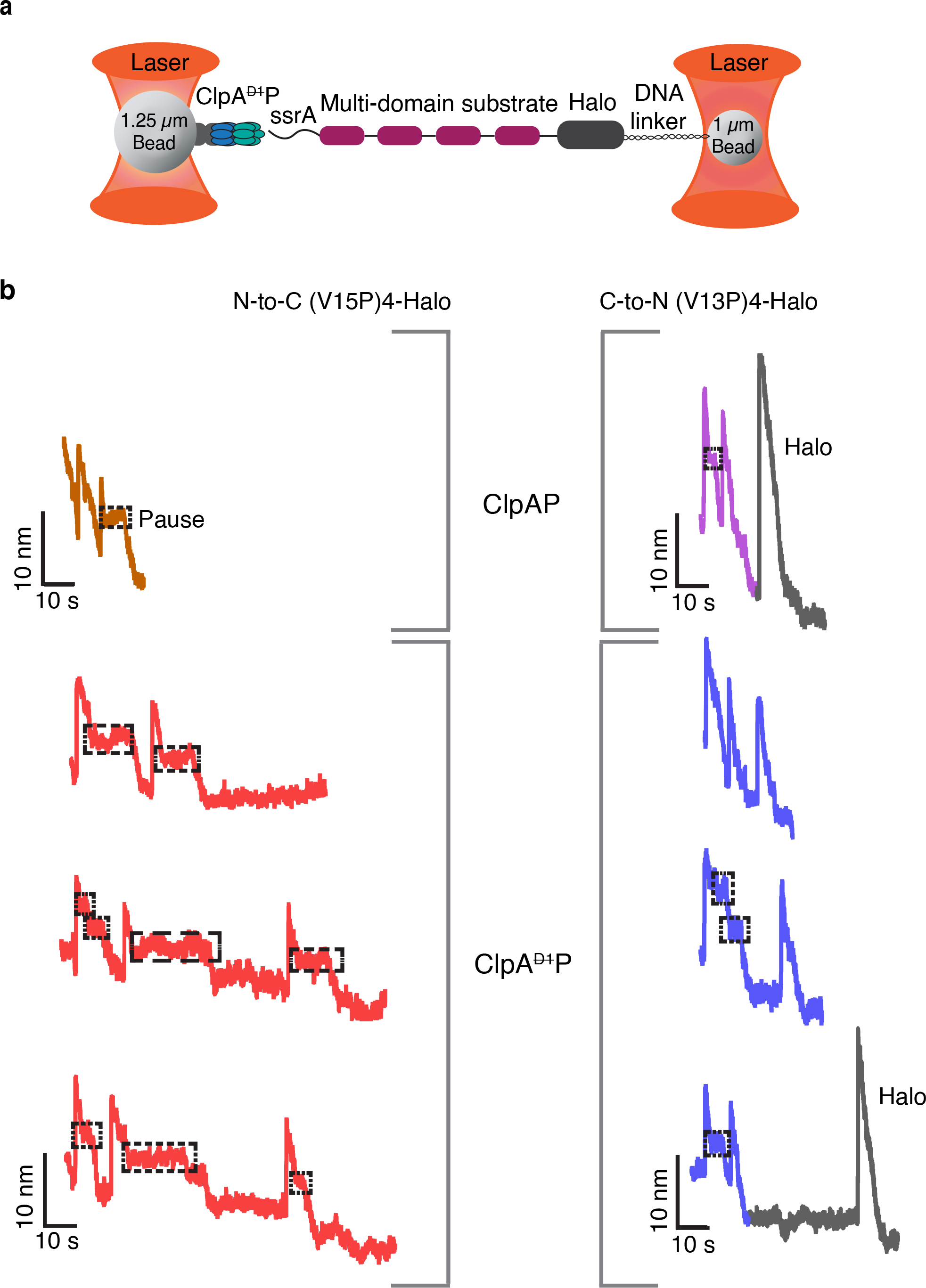
Single-molecule degradation by ClpAP and ClpA^D1^P. (**a**) Optical-trap setup with ClpA^D1^ attached to one bead via ClpP^platform^ and a multi-domain substrate attached to the smaller bead via linkage of the Halo domain to DNA. (**b**) Representative single-molecule degradation traces for ClpAP and ClpA^D1^P in the N-to-C direction (left) or C-to-N direction (right). Different colors are used for unfolding/translocation of titin domains (orange, red, purple, or blue) or the Halo domain (dark gray). Boxes mark pauses. Traces were recorded at forces between 8 and18 pN and were decimated to 300 Hz. ClpAP traces are from Olivares *et al*^15, 16^ and are shown for comparison.

Representative single-molecule traces revealed that ClpA^D1^P takes longer than ClpAP to process one substrate domain in either the N-to-C or the C-to-N direction (Fig. 2b). Processing time is defined as the interval between completion of one translocation event and completion of the next translocation event and thus includes the pre-unfolding dwell, the unfolding event, and the subsequent translocation event. The average processing time for a V15P domain during degradation initiated from the N-terminus was 36 ± 4 s for ClpA^D1^P versus 14 ± 1 s for ClpAP (average ± 1 SEM). For degradation initiated at the C-terminus, these values for the V13P domain were 18 ± 3 s for ClpA^D1^P and 12 ± 2 s for ClpAP.

### D1 plays a major role in unfolding

Pre-unfolding dwell times, the time between the end of translocation of one domain and the start of the unfolding of the next domain, provide information on the relative difficulty of enzymatic unfolding. In experiments in which degradation and thus unfolding of V15P domains was initiated from the N-terminus, the distribution of ClpA^D1^P pre-unfolding dwell times fit well to a sum of two exponentials (*R*^2^ >0.99) with time constants of 20 s (72% amplitude) and 1.4 s (28% amplitude) as compared to a single-exponential time constant of 0.8 s for ClpAP (Fig. 3a; Table 1)^16^. Thus, the majority of V15P domains are unfolded greater than 20-fold more slowly when the D1 domain is hydrolytically inactive. Moreover, unfolding accounted for more than 50% of the processing time for ClpA^D1^P, but less than 10% for ClpAP. Neither ClpA^D1^P nor ClpAP unfolded the Halo domain in the N-to-C direction (Supplementary Fig. S1), as expected from prior studies^16^.

**Table 1:** Unfolding and Translocation parameters with the substrate N-to-C V15P.

|  | N-to-C V15P |  |  |  |  |
| --- | --- | --- | --- | --- | --- |
|  | Pre-unfolding dwell time (s) | Intermediate lifetime (ms) | Average translocation velocity (nm/s) | Pause-free translocation velocity (nm/s) | Step dwell time (s) |
| **ClpA<sup>D1P</sup>** | 20 ± 0.9* (72%),<br>1.4 ± 0.1* (28%)<br>n = 58 | 27.1 ± 1.5*<br>n = 76 | 0.97 ± 0.06*<br>n = 86 | 2.8 ± 0.1*<br>n = 86 | 0.83 ± 0.002* (80%),<br>7.4 ± 0.03* (20%)<br>n = 798 |
| **ClpAP** | 0.8<sup>‡</sup> | 11.5 ± 0.8* | 2.4 ± 0.1<sup>‡</sup> | 2.9 ± 0.2* | 0.55<sup>‡</sup> |

\*Errors are ± 1 SEM
<sup>‡</sup>From Olivares *et al*<sup>16</sup>

**Figure 3:**
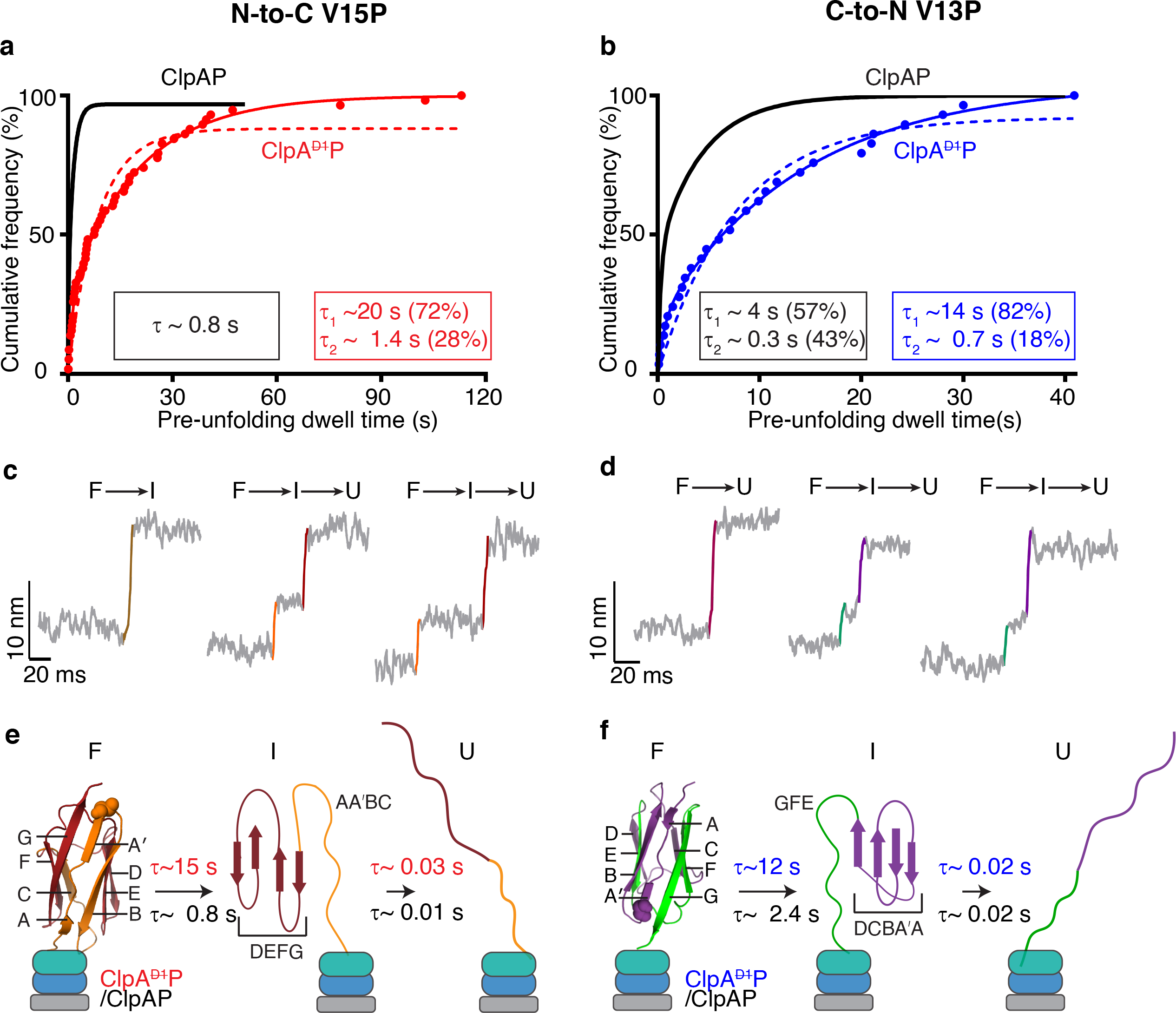
Unfolding by ClpAP and ClpA^D1^P. (**a**) Cumulative-frequency distribution of pre-unfolding dwell times (n=58) for ClpA^D1^P unfolding of V15P domains in the N-to-C direction. The solid red line is a double-exponential fit (*R^2^* > 0.99), with time constants and amplitudes shown in the red rectangle. The red dashed line shows a single-exponential fits. The black line is a single-exponential fit of data for ClpAP unfolding of the same substrate taken from Olivares *et al*.^15, 16^, with the time constant in the black rectangle. (**b**) ClpA^D1^P and ClpAP unfolding of V13P domains in the C-to-N direction. Colors and fits are the same as panel (a), except the ClpAP data is a double-exponential fit. (**c, d**) Traces showing short-lived intermediates during unfolding of titin domains by ClpA^D1^P in the N-to-C (c) or C-to-N (d) directions. Raw traces were decimated to 1500 Hz. (**e, f**) Kinetic and structural models for intermediates formed during enzymatic unfolding of titin domains by ClpA^D1^P and ClpAP. β-strands in the titin domain are labeled, and approximate structural elements in the intermediate during unfolding from each terminus are shown.

During C-to-N degradation, the distribution of ClpA^D1^P pre-unfolding dwell times for V13P also fit well to a sum of two exponentials (*R^2^* > 0.99) with unfolding time constants of 14 s (82% amplitude) and 0.7 s (18% amplitude) as compared to 4.3 s (57% amplitude) and 0.3 s (43% amplitude) for ClpAP (Fig. 3b; Table 2)^15^. Thus, unfolding in the C-to-N direction is ~5-fold slower when D1 is unable to hydrolyze ATP. ClpA^D1^P occasionally unfolded the Halo domain in the C-to-N direction (Fig. 2b), but we obtained too few traces to compile meaningful statistics.

**Table 2:** Unfolding and Translocation parameters with the substrate C-to-N V13P.

|  | C-to-N V13P |  |  |  |  |
| --- | --- | --- | --- | --- | --- |
|  | Pre-unfolding dwell time (s) | Intermediate lifetime (ms) | Average translocation velocity (nm/s) | Pause-free translocation velocity (nm/s) | Step dwell time (s) |
| **ClpA<sup>D1P</sup>** | 14.1 ± 0.9* (82%),<br>0.7 ± 0.1* (18%)<br>n = 29 | 19.0 ± 1.6*<br>n = 38 | 2.1 ± 0.1*<br>n = 44 | 2.9 ± 0.2*<br>n = 44 | 0.6 ± 0.01* (92%),<br>7.4 ± 4.4* (8%)<br>n = 442 |
| **ClpAP** | 4.3<sup>‡</sup> (57%),<br>0.3<sup>‡</sup> (43%) | 19.8 ± 1.9* | 3.0 ± 0.1<sup>‡</sup> | 3.5 ± 0.2* | 0.4<sup>‡</sup> (90%),<br>2<sup>‡</sup> (10%) |

\*Errors are ± 1 SEM
<sup>‡</sup>From Olivares *et al*<sup>15</sup>

Interestingly, ClpAP unfolding of V15P from the N-terminus was faster than the unfolding of V13P from the C-terminus^16, 18^, but this trend was reversed for ClpA^D1^P. Thus, ATP hydrolysis in the D1 domain appears important for aspects of protein unfolding other than simply contributing to the probability of successful unfolding of domains of very high local stability.

### Intermediates in enzymatic I27 unfolding

During V15P and V13P unfolding by ClpA^D1^P, ~30% of events occurred faster than the instrument dead time, indicative of cooperative **F** → **U** unfolding, but ~70% of events displayed non-cooperative denaturation with a short-lived intermediate (**I**). Figures 3c and 3d show examples of non-cooperative unfolding from the N-to-C and C-to-N directions, respectively. Re-analysis of ClpAP unfolding of these domains showed similar trends. The distributions of pre-unfolding dwell times were not altered substantially for two-state and three-state unfolding (Supplementary Fig. S2), indicating that the probability of either class is poorly correlated with the time required for successful unfolding. The **I**-to-**U** lifetimes for C-terminal unfolding by ClpA^D1^P and ClpAP were similar (~20 ms), whereas the **I**-to-**U** lifetime for N-terminal unfolding by ClpA^D1^P (~27 ms) was longer than for ClpAP (~11 ms) (Supplementary Fig. S3a; Tables 1 & 2). Thus, ATP hydrolysis in the D1 ring shortens the intermediate lifetime during unfolding from the N terminus.

Independent of the direction of ClpA^D1^P or ClpAP unfolding, the **F**-to-**I** and **I**-to-**U** distances were ~5 nm and ~9 nm, respectively (Fig. 3c, 3d). These results indicate that the intermediate in N-terminal unfolding is structurally different than the intermediate in C-terminal unfolding. To determine contour lengths for each species, we fit the force dependence of extension using the worm-like-chain (WLC) model and converted these values from nm to the approximate amino acid position in I27, which allowed us to map structural units in the intermediates (Supplementary Fig. S3b). These results suggested that N-terminal unfolding produces an intermediate containing the D, E, F, and G β-strands of I27 (Fig. 3e), whereas C-terminal unfolding resulted in an intermediate containing the A-A’, B, C, and D β-strands (Fig. 3f). Thus, we conclude that ClpAP and ClpA^D1^P, in addition to catalyzing the rate of protein unfolding, alter the unfolding pathways accessible to the I27 domains.

### D1-ring activity is required for efficient translocation

We quantified translocation velocities during degradation by ClpA^D1^P and ClpAP. During N-to-C translocation, the average velocity of ClpA^D1^P (~1 nm/s) was ~60% slower than ClpAP (2.4 nm/s) (Fig. 4a; Table 1)^16^. During C-to-N translocation, the average velocity of ClpA^D1^P (2.1 nm/s) was ~30% slower than ClpAP (3 nm/s) (Fig. 4b; Table 2)^15^. Notably, during ClpA^D1^P translocation, pausing or stalling for significant periods was often observed (see boxed regions in Fig. 2b). By contrast, ClpAP pausing during translocation was infrequent and short lived, as previously observed^15, 16^.

**Figure 4:**
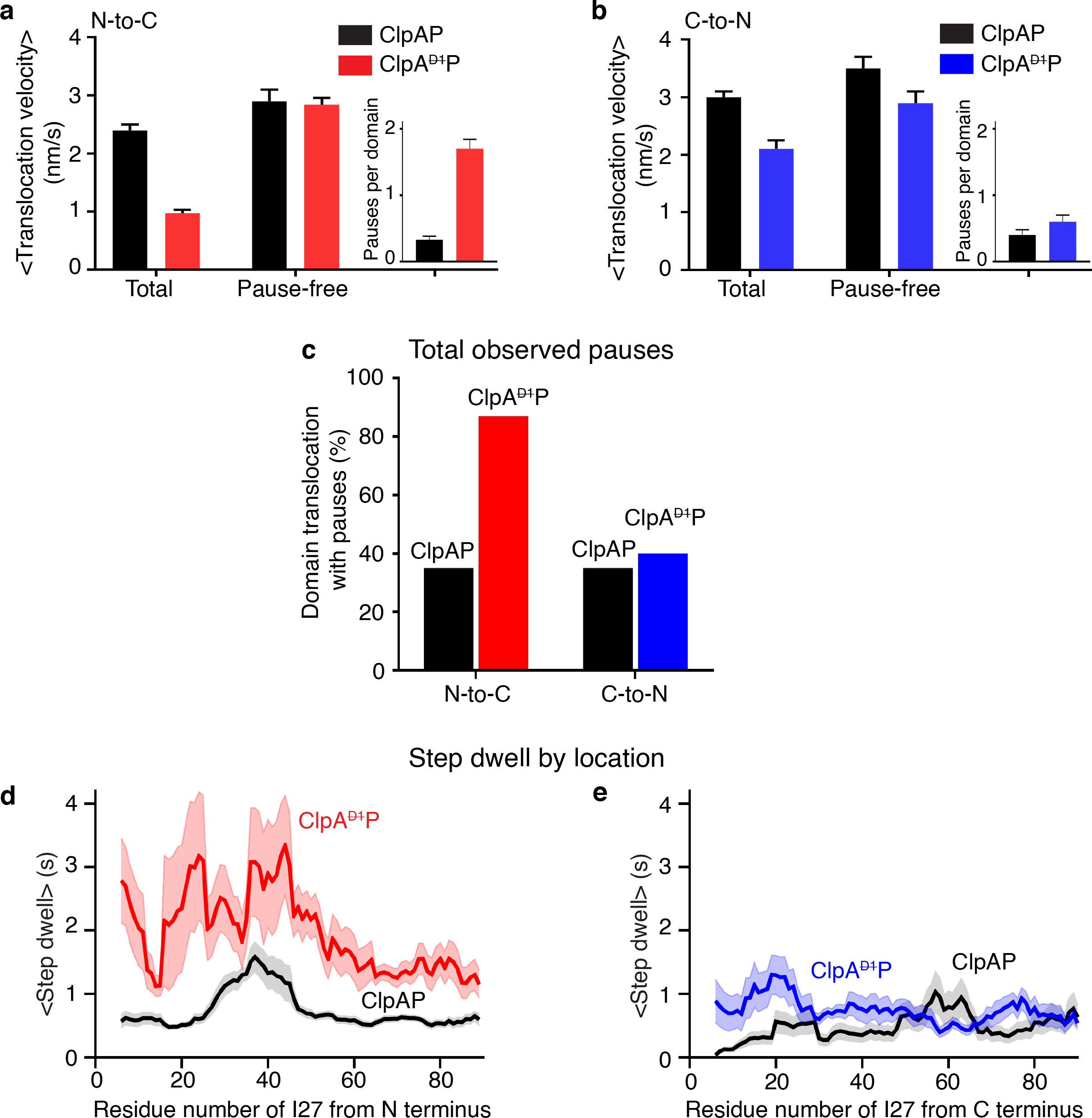
N-to-C and C-to-N translocation. (**a**) Average N-to-C translocation velocities and pause-free velocities for ClpA^D1^P and ClpAP. The inset shows the average number of pauses for one titin domain. Values are the mean ±1 SEM (n=86). **(b)** Average C-to-N translocation velocities and pause-free velocities. Other details are the same as in panel (a). (**c**) Percentage of events with at least one pause during N-to-C or C-to-N translocation of one titin domain for ClpA^D1^P and ClpAP. (**d**) Average step dwell time(s) at each amino acid during N-to-C translocation of a titin domain by ClpA^D1^P (red) or ClpAP (black). The average pre-step dwell time at each amino acid was calculated, and a moving average over a 10-residues window is plotted. The shaded region illustrates ±1 SEM of the moving average at each amino acid. (**e**) Same analysis as in panel (d) except for C-to-N translocation with ClpA^D1^P data colored blue.

When ClpA^D1^P translocation progresses normally, the enzyme takes a step at least every 2.5 s, with most step dwell times being ≤ 1s (Table 1 and 2). We re-calculated average translocation velocities with pauses ≥ 2.5 s removed from each trace. The pause-free velocities for ClpA^D1^P were ~3.0 nm/s for both N-to-C and C-to-N translocation, similar to values determined for ClpAP (Fig. 4a, 4b). These results indicate that the slower translocation velocity of ClpA^D1^P is almost entirely caused by more frequent and longer pauses. Additionally, we observed no force dependence for either the overall or pause-free translocation velocities during this analysis (Supplementary Fig. S4a).

### Directionality influences pause frequency and duration

On average, ClpA^D1^P paused more frequently during N-to-C translocation (1.7 pauses per domain) than in C-to-N translocation (0.55 pauses per domain) (insets in Fig. 4a, 4b), accounting for ~75% and ~30% of the total translocation time, respectively. Furthermore, ~90% of ClpA^D1^P N-to-C translocation events had one or more pauses per domain, whereas this value was ~40% for C-to-N translocation (Fig. 4c). Finally, the longest ClpA^D1^P pauses during N-to-C translocation were ~2 min, whereas this value was ~10 s during C-to-N translocation (see Fig. 5c, 5d). These dramatic differences in pausing frequency and duration, dependent on the direction of movement were not observed for ClpAP translocation. Hence, directional differences in pause frequency and duration for ClpA^D1^P are likely to arise because ATP hydrolysis in the D1 domain alters interactions with the substrate in a manner that depends on the polarity of polypeptide translocation.

**Figure 5:**
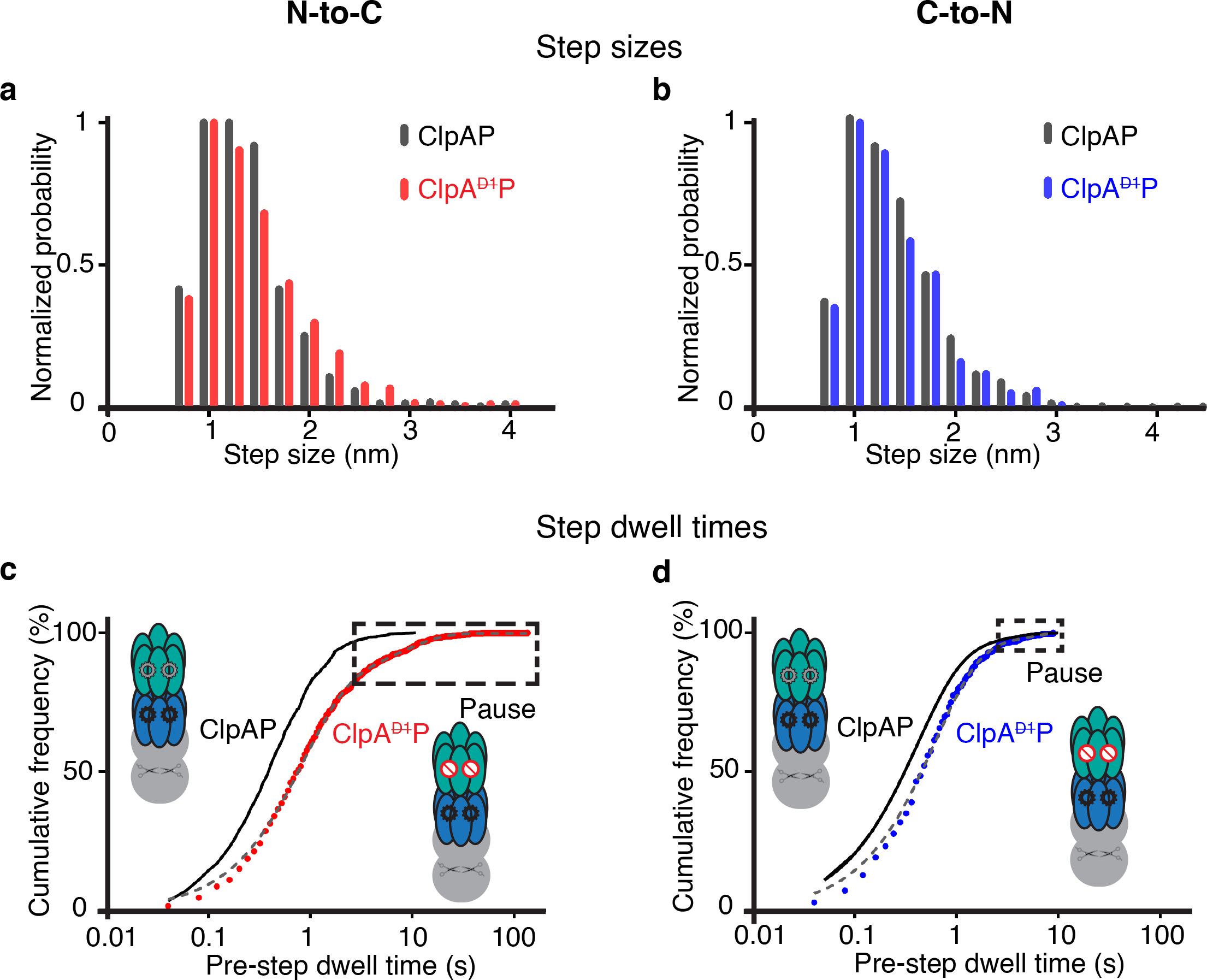
Translocation steps. (**a**) Distribution of step sizes during N-to-C translocation for ClpAP (dark gray) and ClpA^D1^P (red). **(b)** Distribution during C-to-N translocation for ClpAP (dark gray) and ClpA^D1^P (blue). (**c**) Cumulative-frequency distributions of pre-step dwell times during N-to-C translocation by ClpA^D1^P. **(d)** Distributions of pre-step dwell times during ClpA^D1^P C-to-N translocation. In panels (c) and (d) the distributions are fit by double exponentials (gray dashed line, *R^2^* > 0.99). Pre-step dwell times during translocation by ClpAP are shown in black for comparison^15, 16^. Boxes mark pauses, defined as step dwell times ≥ 2.5 s.

To map ClpA^D1^P or ClpAP pause locations, we converted translocation length to amino acid positions using the WLC model and then determined the average dwell time (referred to as the step dwell) at each I27 residue in both translocation directions. The average step dwell at each amino acid for N-to-C translocation was ~1.9 ± 0.7 s for ClpA^D1^P and 0.7 ± 0.3 s for ClpAP, whereas these values were 0.7 ± 0.2 s and ~0.5 ± 0.2 s, respectively, for C-to-N movement (Figs. 4d, 4e). Furthermore, this analysis demonstrated that pausing by ClpA^D1^P occurs throughout the polypeptide sequence, regardless of translocation direction (Supplementary Fig. S4b).

### Step-size distributions and dwell times

In principle, elimination of ATP hydrolysis in the D1 ring might result in shorter translocation steps or longer dwell times between steps. To test these possibilities, we used a stepping algorithm^20^ to analyze translocation-step properties. For both ClpA^D1^P and ClpAP, the step-size distributions were very similar with a peak at ~1 nm, irrespective if translocation was N-to-C or C-to-N (Figs. 5a, 5b). Thus, elimination of ATP hydrolysis in the D1 ring does not alter translocation step size in a significant fashion.

Figs. 5c, 5d show the distribution of pre-step dwell times for ClpA^D1^P and ClpAP. For both translocation directions, dwell times were longer for ClpA^D1^P than for ClpAP, but this difference was greater for N-to-C translocation (Figs 5c, d and Table 1, 2). Thus, inactivation of ATP hydrolysis in the D1 ring lengthens the time between most translocation steps and frequently results in enzyme stalling.

### ATP dependence of ClpA^D1^P activity and substrate slipping

The studies presented above used 5 mM ATP, whereas half-maximal hydrolysis by ClpA^D1^P occurred at ~0.35 mM ATP in the presence of substrate (Supplementary Fig. S5a). Prior studies show that the K_M_ of ATP for the D1 ring is tighter than that for the D2 ring^14^. To determine the ATP dependence of ClpA^D1^P pausing behavior during unfolding and translocation, which should reflect D2-ring activity, we also performed single-molecule N-to-C degradation assays at 0.1 mM (n = 18), and 0.25 mM (n = 37) ATP, concentrations where the D2 ring would not be fully ATP bound. Notably, the distribution of pre-unfolding dwell times and translocation velocities were not substantially altered at these lower ATP concentrations (Fig. 6a, 6b). We conclude that full saturation of the D2 ring with ATP is not required for robust mechanical function.

**Figure 6:**
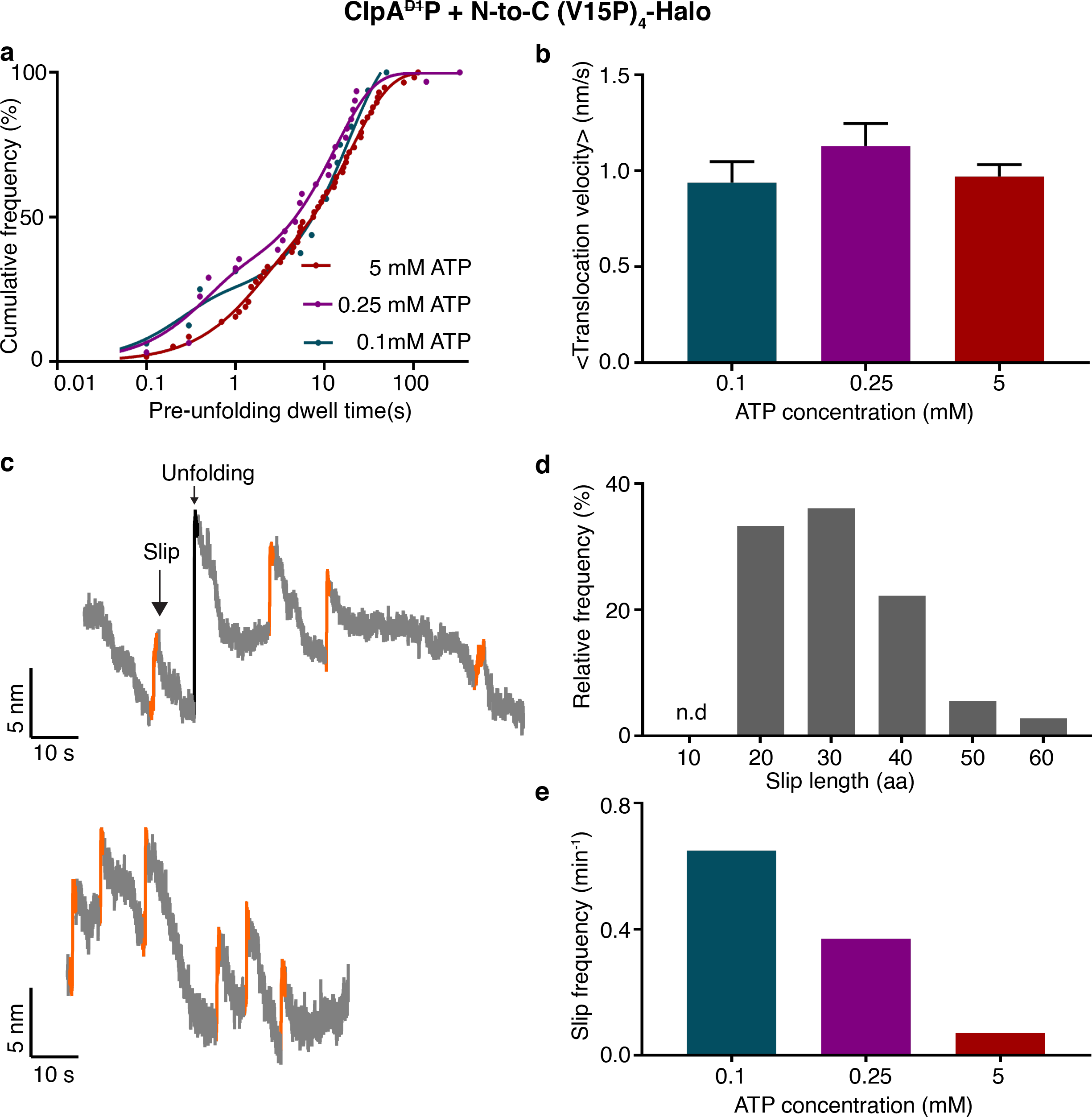
ATP dependence of ClpA^D1^P degradation. (**a**) Distribution of pre-unfolding dwell times for N-to-C degradation at different ATP concentrations. The solid lines are double-exponential fits. Time constants are 20 ± 0.9 s and 1.4 ± 0.1 s for 5 mM ATP; 13 ± 1.5 s and 0.4 ± 0.1 s for 0.25 mM ATP; and 19 ± 4 s and 0.2 ± 0.1s for 0.1 mM ATP. (Values are average ± 1 SEM). (**b**) Average translocation velocities at different ATP concentrations. (Values are mean ± 1 SEM). (**c**) Representative traces (decimated to 300 Hz) showing slipping events (orange) during N-to-C unfolding and translocation by ClpA^D1^P using 0.1 mM ATP. (**d**) Relative frequency of substrate slips of different lengths (number of amino acids). (**e**) Total substrate-slip frequencies decrease as the ATP concentration increases.

In ensemble experiments, by contrast, ClpA^D1^P degradation of the multidomain substrate in the N-to-C direction and GFP-ssrA in the C-to-N direction were both slowed at ATP concentrations below 0.75 mM (Supplementary Fig. S5b, S5c). Moreover, steady-state kinetic parameters determined for ClpA^D1^P degradation of GFP-ssrA at different ATP concentrations revealed a substantial V_max_ decrease at 0.1, 0.25 and 0.5 mM ATP (Supplementary Fig S5e). The differences observed between single-molecule and ensemble degradation at low ATP concentrations are likely to reflect inefficient substrate engagement by ClpA^D1^P, a step that is not monitored in the optical-trap assay.

Suboptimal ATP concentrations did, however, alter one important feature of the mechanical processes of unfolding and translocation. During N-to-C single-molecule degradation by ClpA^D1^P, we observed periodic “slips”, in which the substrate polypeptide appeared to thread backwards through the enzyme, causing a fast increase in bead-to-bead distance (Fig. 6c). In contrast to the changes in bead-to-bead distance associated with unfolding, slipping resulted in variable distance changes that were shorter than the I27-contour length and also often occurred before domain translocation was complete. Most slips corresponded to ~20-40 amino acids (Fig. 6d). Slipping was more common at lower ATP concentrations with a total frequency of 0.65 slips/min at 0.1 mM ATP, 0.37 slips/min at 0.25 mM ATP, and 0.07 slips/min at 5 mM ATP (Fig. 6e). Substrate slipping was infrequent for ClpAP, even at low ATP concentrations. In combination, these observations suggest that ClpA^D1^P has a decreased ability to grip substrates properly, and this feature is unmasked by lower ATP concentrations where the active D2 ring is unlikely to be saturated with ATP.

## Discussion

Protein unfolding and remodeling enzymes that contain two AAA+ rings play important roles in proteostasis in all cells, but no clear or unifying function for the second motor ring has emerged^6^. Prior studies show that the D2 ring of ClpAP plays the dominant role in ATP-dependent degradation^14^. Our optical-tweezers studies here probe the consequences of eliminating ATP hydrolysis in the D1 ring of ClpA on single-molecule unfolding and translocation by the ClpA^D1^P protease. Similarities between ClpA^D1^P and ClpAP reveal ring functions specific to ATP hydrolysis in the D2 ring, whereas differences provide a view into mechanistic tasks that normally require ATP hydrolysis in the D1 ring.

Compared to ClpAP, ClpA^D1^P is a slower protein translocase because it pauses more frequently during polypeptide translocation, a defect that is more severe during N-to-C than C-to-N translocation. Likewise, compared to ClpAP, the average time required for ClpA^D1^P to unfold the V15P domain is ~20-times slower in the N-to-C direction, whereas unfolding of the V13P domain is ~5-times slower in the C-to-N direction. Thus, ClpA^D1^P is both a less reliable protein translocase and a slower protein unfoldase. These mechanical defects are likely to be related, as slower ClpA^D1^P unfolding and increased pausing are both intensified during N-to-C degradation; “D2 motor stalling” can reasonably explain both effects and would be expected to be more severe against the resisting force that occurs during unfolding attempts.

Prior studies of a ClpA^D1^P variant show that it degrades substrates of low and intermediate stability as efficiently as ClpAP, whereas degradation of high-stability substrates required ATP hydrolysis in both the D1 and D2 rings^14^. Thermodynamically and kinetically, however, the domains studied here are metastable (V13P – ΔG_u_ = 2.9 kcal/mol, *k*_u_ = 32 min^−1^: V15P – ΔG_u_ =4.6 kcal/mol; *k*_u_ = 2.3 min^−1^)^21^ and yet are unfolded ~5 to ~20-times more slowly by ClpA^D1^P than by ClpAP. Thus, our results indicate that ATP hydrolysis in both the D1 and D2 rings can be important for enhancing unfolding of protein domains of moderate and well as high stabilities.

Although ClpAP and ClpA^D1^P catalyze similar rates of ATP hydrolysis in the absence of protein substrates as shown here and previously^14^, ClpA^D1^P hydrolyzes ATP at only 50-60% of the wild-type rate in the presence of either of our protein substrates. This change in ATPase rate, at least in part, appears to be a consequence of two interrelated factors: (*i*) ClpA^D1^P spends more of the total time during degradation carrying out unfolding of substrate domains than does ClpAP; and (*ii*) ATP hydrolysis by AAA+ machines is slower during unfolding than translocation^21^.

Regardless of direction, the majority of I27-unfolding events catalyzed by ClpAP or ClpA^D1^P occur via an intermediate. The intermediate is different during N-to-C than C-to-N unfolding, and is also longer lived during N-to-C unfolding by ClpA^D1^P. By contrast, non-enzymatic I27 unfolding is generally cooperative or two-state; and rare intermediates when observed, are clearly distinct from those characterized here^22–24^. These differences highlight the fact that AAA+ enzymes, like ClpAP change protein-unfolding pathways by pulling directly on one end of the target molecule, a very different process than spontaneous or force/chemical-induced denaturation.

During N-to-C or C-to-N translocation, the step sizes taken by ClpAP and ClpA^D1^P are indistinguishable (average ~1.25 nm) and the pause-free velocities are similar (~3 nm/s). Thus, motor activity in the D2-ring alone is sufficient for normal step sizes and near normal translocation rates when pausing is ignored. In the absence of ATP hydrolysis in the D1 ring, however, pausing slowed the overall rate of translocation more than 2-fold in the N-to-C direction and ~30% in the C-to-N direction. Defects in ClpA^D1^P unfolding are also likely to be caused by pausing or stalling of the D2 motor.

Compared to wild-type ClpAP translocation and unfolding, the ClpA^D1^P defects are more severe in the N-to-C direction as shown by: *(i)* longer pre-unfolding dwell times; *(ii)* higher frequencies of translocation pauses; *(iii)* longer pause durations; and *(iv)* slightly longer step dwells. Thus, the wild-type D1 of ClpA ring facilitates more efficient mechanical processing of substrate proteins, especially in the N-to-C direction. Previous instances of AAA+ proteases degrading substrates more efficiently from one terminus have been typically ascribed to the substrate structure, such that unfolding of a given protein is “easier” from one end and thus the rates of degradation from the “easy” terminus are faster than those from the “difficult” end^14, 16, 25^. Our results, however, show substantial D1-dependent differences in translocation (in addition to the unfolding effects) in the two directions using the same protein substrates. We propose that interactions between the D1 ring and substrate polypeptide are more favorable for the N-to-C polarity because of stereo-chemical differences in enzyme-substrate contacts. Perhaps the D2 ring has the opposite-polarity preference due to a differently shaped enzyme-substrate gripping surface, thus explaining why ClpA^D1^P with an inactive D1 motor has a harder time carrying out mechanical processes when traveling N-to-C on a polypeptide.

How does an active D1 ring suppress pausing? Because ClpA^D2^P (ATPase defective D2 ring) degrades some target proteins slowly^14^, and the D1 ring contains AAA+ motifs required for motor activity, it is likely that the wild-type D1 ring can support translocation through the ClpA pore and into ClpP. However, ClpA^D2^P hydrolyzes ATP at least 10-fold more slowly than either ClpAP or ClpA^D1^P^14^, likely accounting for its strong defect in degradation. Together, these observations establish that D2 is the principal unfoldase/translocase engine, whereas D1 serves as an auxiliary motor. Why then is the slow and inefficient D1 motor required for robust translocation and unfolding by wild-type ClpAP? One possibility is that the D1 and D2 motors function independently. Thus, when the dominant D2 motor fails, the slower D1 motor fills in and prevents pausing. Independence is supported by the fact that the contributions of the D1 and D2 rings to overall ATP hydrolysis are roughly additive under some conditions^14^. Moreover, the greater importance of the D1 ring during N-to-C than C-to-N translocation and unfolding is also consistent with an independent model. A second and non-exclusive possibility is that the two motor rings are allosterically coupled, as proposed for Hsp104^26^. For example, allosteric coupling might allow ATP hydrolysis in the D1 ring to restart the D2 motor when it stalls.

The D1 ring appears to play an additional important role in gripping protein substrates. For example, ClpAP typically unfolds substrates faster but translocates them more slowly than ClpXP, which has just one AAA+ ring^15^. Faster unfolding by ClpAP would be expected if it generates a similar unfolding force but can grip substrates more tightly using two rings compared to the single ring of ClpXP. Within the pores of AAA+ unfoldases there are highly conserved pore-1-loops (consensus sequence GYVG) that are especially important for substrate binding within, and translocation through the unfoldase. In support of the idea that the larger AAA+ enzymes may grip substrates better throughout their pores, cryo-EM structures of the double-ring Hsp104 enzyme reveal from 7 to 10 contacts between pore-1-loops and substrate, whereas only 4 or 5 equivalent contacts are observed in structures of single-ring enzymes^27, 28^. Our experiments here strengthen the model that D1 and D2 both contribute to substrate grip, and that different features of each ring provide a better grasp of its target. At low concentrations of ATP, where the D2 ring is expected to be incompletely nucleotide bound (its *K_M_* for ATP hydrolysis is ~13-fold weaker than that of D1^14^), we observe polypeptide “back slipping” through the enzyme pore of ClpA^D1^P but not ClpAP. Thus, ATP hydrolysis in the D1 ring of wild-type ClpAP appears to help the enzyme maintain a grip on the substrate and prevent back-slipping when D2 is not fully saturated with ATP. Interestingly, compared to ClpXP, ClpAP plays a more substantial role in degradation during stationary phase^29^, a condition where ATP concentrations can drop considerably^30^.

Remarkably, like ClpA, other well-studied double-ring AAA+ unfoldases/remodeling enzymes have one ring that is the dominant and fast ATPase, whereas the other ring is much slower^31^. In most cases, as with ClpA, it is the D2 ring that is dominant, although this architecture appears to be flipped for yeast Hsp104^32^. In two cases (Hsp104 and Cdc48), the faster ATPase is thought to be the major translocation motor, as with ClpA, and emerging evidence favors models where the auxiliary ring has an important role in substrate release from the enzyme^33, 34^. Interestingly, the two rings in ClpAP evolved from different subfamilies of AAA+ unfoldases^35^, rather than duplication of the AAA+ module of a single-ring enzyme, further supporting the view that each ring performs unique functions in the double-motor enzyme. The D1 ring of ClpA is a member of the classic clade of AAA+ enzyme^35^. Some members of this clade, including YME1, PAN, and Rpt_1-6_, are thought to operate by strictly sequential ATP-hydrolysis mechanisms that result in two-residue translocation steps^28, 36, 37^. The D2 of ClpA ring, is a member of the HCLR clade, together with ClpX, HslU, and Lon^35^. ClpX and HslU appear to operate by probabilistic mechanisms of ATP hydrolysis, and ClpX takes basic translocation steps of ~6 residues, similar to those of ClpAP’s and ClpA^D1^P’s translocation mechanism^17, 38, 39^. It is straightforward to picture situations in which having an engine of each type in a two-motor machine might be advantageous. Further dissection of individual AAA+ enzymes and their rings should reveal how differences in mechanism support the various biological functions of this large and diverse family of protein unfoldases and remodeling machines.

## Materials and Methods

### Protein Purification

ClpA^D1^, a variant of *E. coli* ClpA – containing the ΔC9 deletion^40^, which prevents autodegradation without affecting activity, and E286Q Walker-B mutation – was cloned, expressed and purified as described^15, 16, 41^. Briefly, a pET23b plasmid carrying the gene encoding ClpA^D1^ was transformed into a BL21(DE3) strain with the wild-type *clpA* gene disrupted. Cells were grown to OD_600_ ~1 at 30 °C, induced with 0.4 mM IPTG, grown for additional 2 h, and harvested. The cell paste was resuspended in Lysis buffer (50 mM Tris-Cl, pH 7.5, 2 mM EDTA, 10% glycerol, 2 mM DTT) and stored at –80 °C. After lysis by French press, the lysate was clarified by centrifugation at 30,000 x g for 30 min, and 40% (w/v) ammonium sulfate was added to precipitate ClpA^D1^. The pellet was resuspended in SBA buffer (25 mM HEPES, pH 7.5, 200 mM KCl, 0.1 mM EDTA, 10% glycerol, 2 mM DTT), and ClpA^D1^ was purified by ion-exchange chromatography using S-Sepharose and MonoQ HR 5/5 columns (GE Healthcare). Purified ClpA^D1^ was dialyzed into storage buffer (50 mM HEPES, pH 7.5, 20 mM MgCl_2_, 0.3 M NaCl, 300 mM arginine, 10% glycerol, 0.5 mM DTT), divided into aliquots, and stored at –80 °C. ClpA^ΔC9^ was used as the wild-type background for ClpA^D1^ here and previously^15^. ClpA^ΔC9^, ClpP, and ClpP^platform^ (used for optical-trap experiments) were expressed, and purified as described^15, 42^, as were the Halo-(V13P)_4_-ssrA substrate (C-to-N)^15, 18^ and Cys-(V15P)_4_-Halo substrate (N-to-C)^16^. An ssrA peptide containing an N-terminal maleimide was subsequently attached to the N-terminus of Cys-(V15P)_4_-Halo to generate ssrA-(V15P)_4_-Halo^16^.

### Single-molecule optical-trapping

For optical-trap experiments, ClpA^D1^/ClpP^platform^ was attached to one bead and a titin-ssrA substrate was attached to the other bead as described^15, 17-19^. Briefly, one end of a biotinylated DNA linker (3500 base pairs) was attached to a 1.09 μm streptavidin-coated polystyrene bead (Spherotech) on the other end was attached to Halo-domain of one of the multidomain substrates. The 1.09 μm bead was tethered to a glass coverslip using a DNA-linked glass-binding peptide aptamer. Biotinylated ClpP^platform^ was attached to a 1.25 μm streptavidin-coated polystyrene bead (Spherotech), and ClpA^D1^ was added in presence of 5 mM ATP (except for analysis of ATP-concentration effects). After complex formation, free ClpA^D1^ was removed by centrifugation and washing of the beads. Experiments were performed at 17-20 °C in PD-T buffer (25 mM HEPES, pH 7.6, 100 mM KCl, 10 mM MgCl_2_, 10% glycerol, 0.1% Tween-20, and 1 mM Tris-2-carboxyethyl-phosphine) containing ATP-regeneration and oxygen-scavenging systems as well as 1 mg/mL bovine serum albumin (Sigma-Aldrich). The beads were trapped by two 1064-nm lasers in a passive-force clamp mode, and tethers between ClpA^D1^P and the substrate were formed by moving the weakly-trapped larger bead.

### Data Analysis

Data acquisition was carried out as described^15, 18^. Custom MATLAB scripts were used to calculate the inter-bead distance, unfolding distance, pre-unfolding dwell times, and to determine step sizes and step-dwell times. Average translocation velocities were determined by dividing the sum of step sizes by the sum of step dwells for translocation including pauses for individual domains. Data were collected at 3 kHz and were decimated to 50 Hz, and individual steps were determined using χ^2^ minimization method as described^20^. Steps < 0.75 nm and backward steps were combined, and the dwell time preceding a combined step was added to the dwell time of the following step. Any step dwell ≥ 2.5 s was considered a pause and removed from the total step-dwell time when calculating pause-free translocation velocities.

## Acknowledgments

We thank Adrian Olivares, Kristin Zuromski, Sora Kim, and Gina Mawla for materials, advice, and helpful discussions. This work was supported by NIH Grants GM-101988 and AI-15706 and the Howard Hughes Medical Institute. H.C.K and T.A.B. are employees of the Howard Hughes Medical Institute.

## Author Contributions

H.C.K., R.T.S., and T.A.B. designed the research; H.C.K. performed the research; H.C.K., R.T.S., and T.A.B. analyzed the data; and H.C.K., R.T.S., T.A.B. wrote the paper.

Authors declare no competing interests.

