## Supplementary Information for "The non-dominant AAA+ ring in the ClpAP protease functions as an anti-stalling motor to accelerate protein unfolding and translocation"

**Supplementary Methods:**

*Biochemical Assays:*

*ATP hydrolysis:* Steady-state rates of ATP hydrolysis were measured at 24 °C in an NADH-coupled assay by monitoring loss of absorbance at 340 nm (1) using 100 nM $ClpA_6$ or $ClpA^{D1}{}_6$, 200 nM $ClpP_{14}$, and 5 µM substrate. The final reaction mixtures were in PD-T buffer supplemented with 5 mM ATP (unless noted in legends), 0.7 mM NADH, 6.25 mM phosphoenolpyruvate, 23.5 U/mL pyruvate kinase, and 20 U/mL lactate dehydrogenase.

*Ensemble protein degradation:* The kinetics of degradation of GFP-ssrA by $ClpA^{D1}{}_6$ (0.6 µM) and $ClpP_{14}$ (1.6 µM) were assayed by monitoring loss of GFP fluorescence at 540 nm (excitation 470 nm) at 24 °C in PD-T buffer.

The ssrA-$(V15P)_4$-Halo substrate (100 µL; 1 µM) was fluorescently labeled by incubation with HaloTag TMR ligand (0.5 µL; 0.5 mM; Promega) in 25 mM HEPES (pH 7.5), 100 mM KCl, 10 mM $MgCl_2$, 10% glycerol, 0.1% Tween-20, and 1 mM DTT at 30 °C for 15 min. Degradation reactions at room temperature contained $ClpA_6$ or $ClpA^{D1}{}_6$ (200 nM), $ClpP_{14}$ (400 nM), ATP (0.1 to 5 mM), and an ATP-regeneration system. Samples were taken at different time points, and degradation was quenched by addition of SDS-sample buffer and flash freezing in liquid $N_2$. Samples were thawed, boiled for 5 min, and a fraction of each reaction was electrophoresed on a Mini-PROTEAN TGX 4–15% (w/v) precast gel (Bio-Rad) prior to scanning the gel using a Typhoon FLA 9500 scanner (GE Healthcare).

*Data Analysis:*

*Intermediate structures in titin[I27] unfolding:* The extensions for F→I, I→U, and F→U transitions were arranged in increasing order of experimental force, and a moving average of three data points was fit to the WLC model (persistence length 0.36 nm) to determine the contour length (Lc) associated with each transition. The contour length change (ΔLc) was given by the equation, $\Delta Lc = Lc - (d_f - d_r)$, where $d_f$ is the length of titin[I27] folded domain (PDB 1TIT) and $d_r$ is the length the remaining folded portion. ΔLc was converted into number of amino-acid residues using a value of 0.365 nm/residue and mapped onto terminal structural units of the titin[I27] domain.

**Table S1: Thermodynamic, kinetic, mechanical stabilities of common degradation substrates.**

| Substrate | $\Delta G_u$ (kcal mol$^{-1}$) | $k_u$ (min$^{-1}$) | $F_u$ (pN) [*pulling speed nm/s*] | Reference |
|---|---|---|---|---|
| V13P titin | 2.9 | 32 | 132 [*600*] | (2) |
| V15P titin | 4.6 | 2.3 | 159 [*600*] | (2) |
| λ-repressor N domain | 4.8 | 300 | not determined | (3, 4) |
| T4 lysozyme | 14.1 | 0.4 | 50 [*400*] | (5-7) |
| GFP | 8 | $10^{-9}$ | 104 [*300*] | (2, 8) |

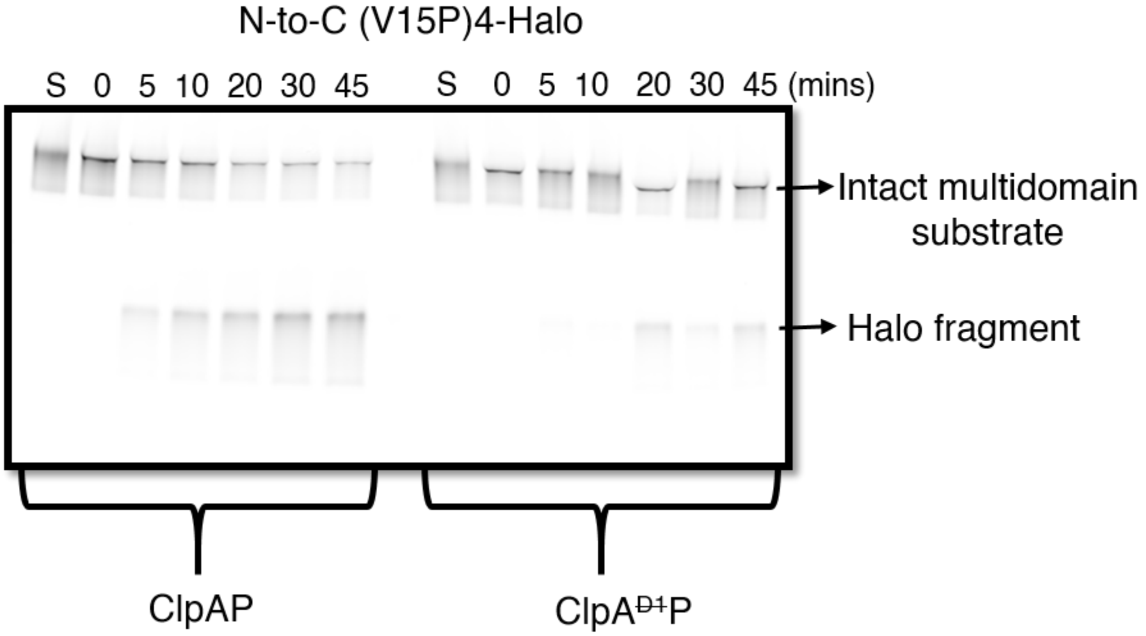


**Figure S1: Gel assay of substrate degradation by ClpAP and ClpA$^{D1}$P.** SDS PAGE assay of degradation of fluorescent ssrA-(V15P)$_4$-Halo by ClpAP and ClpA$^{D1}$P. Degradation from the N-terminus results in accumulation of a fluorescent fragment corresponding to the TMR-labeled C-terminal Halo domain.

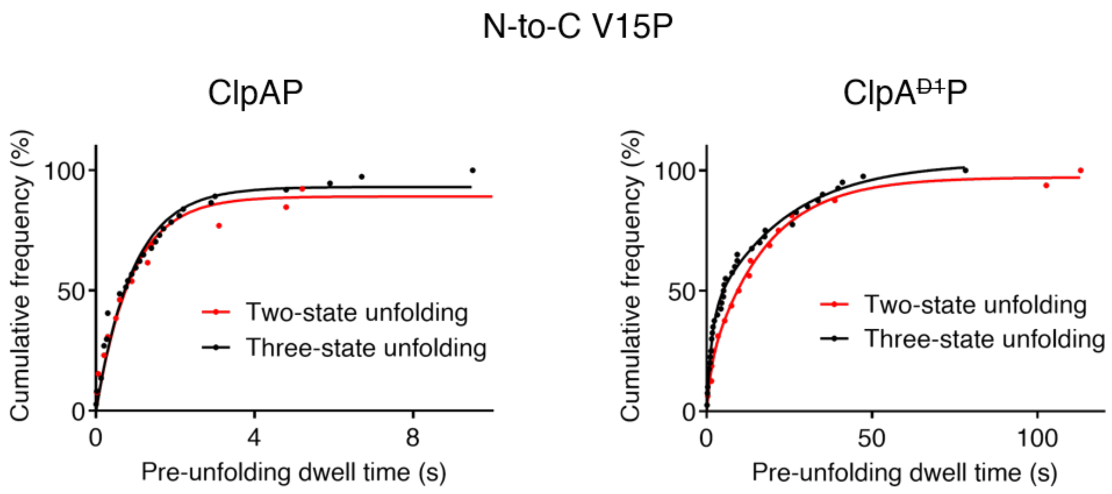


**Figure S2: Pre-unfolding dwell times for two-state and three-state denaturation.** Cumulative-frequency distributions of pre-unfolding dwell times for N-to-C unfolding of V15P titin domains by ClpAP (left) and ClpA$^{Đ1}$P (right). Two-state (red) and three-state (black) unfolding events are plotted separately. For ClpAP, the two-state ($R^2$ = 0.96) and three-state ($R^2$ = 0.97) distributions were fit to single-exponentials ($\tau$ ~0.9 s in both cases). Hence, the probability of ClpAP unfolding by two-state or three-state pathways is unrelated to the time required for initial unfolding. For ClpA$^{Đ1}$P, distributions were fit to double exponentials; fitted $\tau$ values (amplitudes) were 1.2 s (17%) and ~17 s (83%) for two-state unfolding ($R^2$ = 0.99), and 1.5 s (37.5%) and ~23 s (62.5%) for three-state unfolding ($R^2$ = 0.99). Thus, titin V15P domains that unfold via an intermediate are slightly more susceptible to enzymatic unfolding by ClpA$^{Đ1}$P.

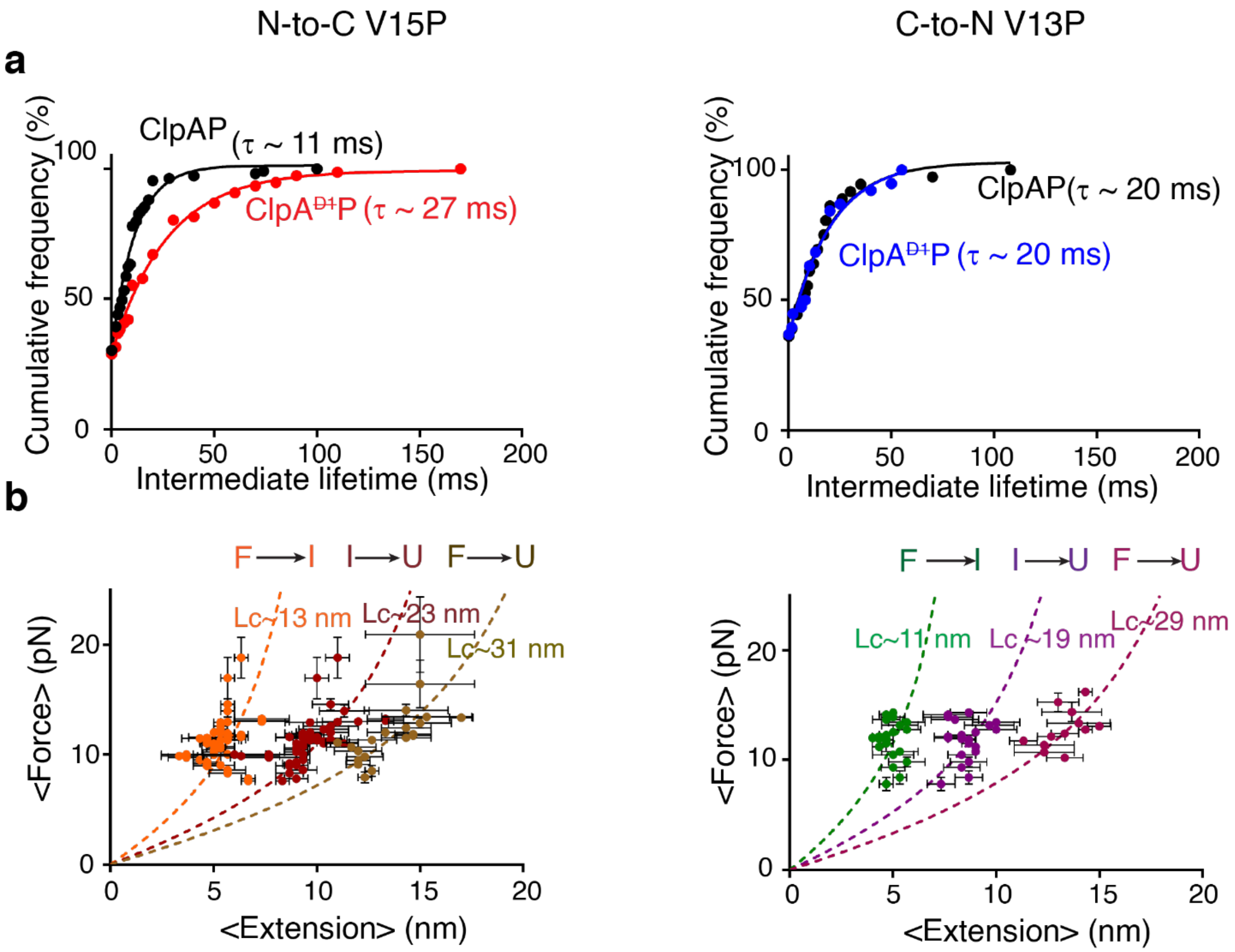


**Figure S3: Intermediate lifetimes and contour lengths. (a)** Cumulative-frequency distribution of lifetimes of the intermediate state during N-to-C (red; left) and C-to-N (blue; right) unfolding by ClpA$^{\text{Ð1}}$P. ClpAP distributions are shown in black. Single-exponential fits are shown as solid lines. **(b)** Force versus extension plots for ClpA$^{\text{Ð1}}$P catalyzed F→I, I→U, and F→U unfolding transitions. The data are means (± 1 SD) of a moving-window average of three points arranged in order of increasing force. Data were fit to the WLC model of polymer elasticity (dotted lines) to obtain Lc contour lengths. Lc values for N-to-C unfolding of V15P were 13.1 ± 0.3 nm (F→I), 23.2 ± 0.4 nm (I→U), and 30.6 ± 0.7 nm (F→U). For C-to-N unfolding of V13P, these values were 11.2 ± 0.3 nm (F→I), 19.4 ± 0.4 nm (I→U), and 28.6 ± 0.6 nm (F→U).

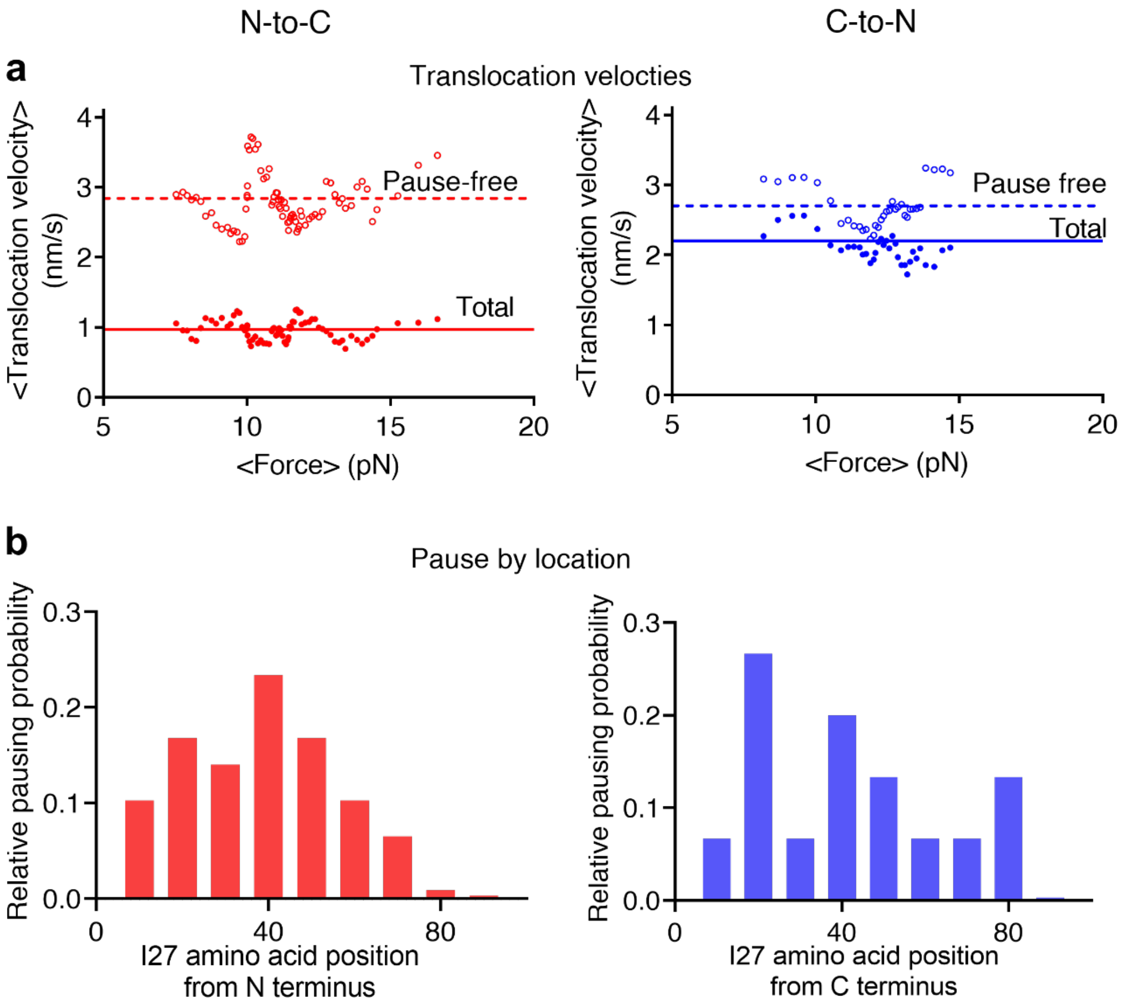


**Figure S4: Translocation as a function of force and pausing locations. (a)** Translocation velocities (closed circles) and pause-free translocation velocities (open circles) for ClpA$^{\text{Ð1}}$P in the N-to-C I27 direction (red; averages 0.96 and 2.8 nm/s, respectively) and the C-to-N direction (blue; averages 2.1 and 2.7 nm/s, respectively). Data plotted are a moving-average of 10 consecutive values ranked by force. These results indicate that translocation velocities are relatively independent of trap force. **(b)** Relative pausing probability of ClpA$^{\text{Ð1}}$P during N-to-C (red) and C-to-N (blue) translocation. Pausing frequency was binned by 10 amino acids and normalized to total pausing probability to calculate relative probability. The distribution observed indicates that enzyme pausing in either direction occurs throughout the titin$^{I27}$ domain sequence.

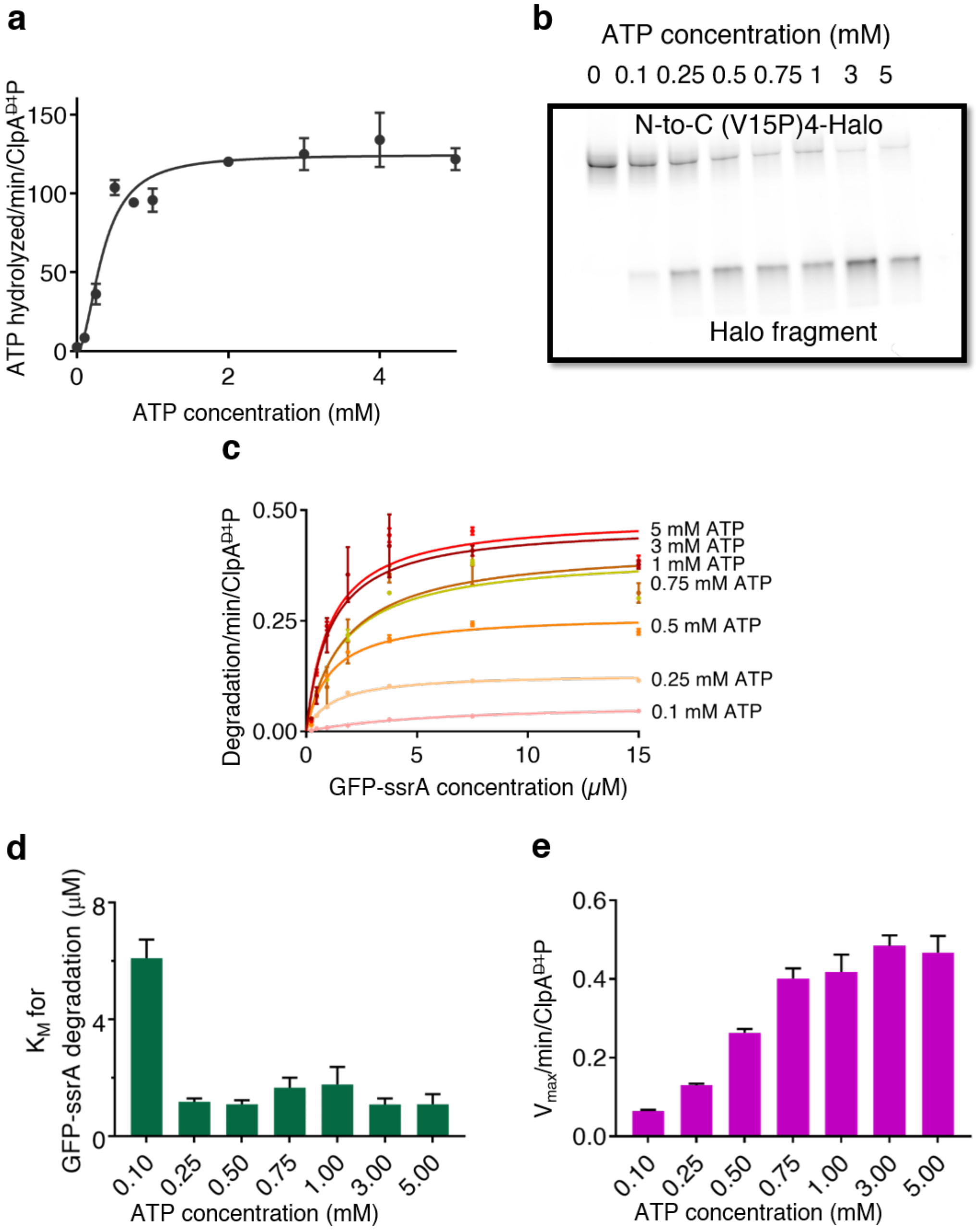


**Figure S5: ATP-concentration dependence of ClpA$^{Ð1}$P activities. (a)** Rate of ATP hydrolysis by ClpA$^{Ð1}$P in the presence of ssrA-(V15P)$_4$-Halo (5 μM) and different ATP concentrations. The half-maximal rate of ATP hydrolysis occurred at ~0.35 mM ATP. **(b)** ATP dependence of ClpA$^{Ð1}$P degradation of fluorescent ssrA-(V15P)$_4$-Halo (1 μM) after 30 min assayed by SDS PAGE. **(c)** ATP-dependence of the rates of ClpA$^{Ð1}$P degradation of different concentrations of GFP-ssrA. Solid lines are fits to the Michaelis-Menten equation. Values are average ± 1 SD of three independent measurements. **(d)** $K_M$ for ClpA$^{Ð1}$P degradation of GFP-ssrA are not significantly altered at lower ATP concentration except at 0.1 mM ATP. **(e)** $V_{max}$ for ClpA$^{Ð1}$P degradation of GFP-ssrA is slower at low ATP concentrations. The error bars are standard errors of the Michaelis-Menten fit.
